# The Microbiome Modeling Toolbox: from microbial interactions to personalized microbial communities

**DOI:** 10.1101/318485

**Authors:** Federico Baldini, Almut Heinken, Laurent Heirendt, Stefania Magnusdottir, Ronan M.T. Fleming, Ines Thiele

## Abstract

**Motivation:** The application of constraint-based modeling to functionally analyze metagenomic data has been limited so far, partially due to the absence of suitable toolboxes.

**Results:** To address this shortage, we created a comprehensive toolbox to model i) microbe-microbe and host-microbe metabolic interactions, and ii) microbial communities using microbial genome-scale metabolic reconstructions and metagenomic data. The Microbiome Modeling Toolbox extends the functionality of the COBRA Toolbox.

**Availability:** The Microbiome Modeling Toolbox and the tutorials at https://git.io/microbiomeModelingToolbox.

## 1 Introduction

Microbial community sequencing data are increasingly available for numerous environmental niches, including the human gut. The analysis of this data often relies on investigating which microbes are present in a given sample. However, to further our understanding of the functional contribution of individual microbes in a community as well as the overall functional differences between communities, advanced analysis approaches, such as computational modeling, are required.

One possible approach is the constraint-based reconstruction and analysis (COBRA) approach, which builds genome-scale reconstructions of an organism and enables the prediction of, e.g., phenotypic properties [7]. Through the application of condition-specific constraints, an organism’s metabolic reconstruction can be converted into many distinct condition-specific models. These models can be analyzed using available toolboxes, such as the widely used, Matlab (Mathworks, Inc.) based COBRA Toolbox [4]. Metabolic reconstructions have been assembled for many organisms, including over 800 gut microbes (AGORA) [6] and human [1].

While the COBRA Toolbox encapsulates many tools developed by the community for biotechnological and biomedical applications, it is currently focused on modeling single organisms or cells. Here, we present the Microbiome Modeling Toolbox, a COBRA Toolbox extension, which allows for the assembly of multispecies models, starting from a set of individual metabolic reconstructions, to model microbe-microbe and host-microbe interactions as well as functional properties of microbial communities (Figure 1). By integrating sample-specific metagenomic data, the Microbiome Modeling Toolbox facilitates its analysis in the context of well-defined microbial metabolic reconstructions.

**Figure 1:**
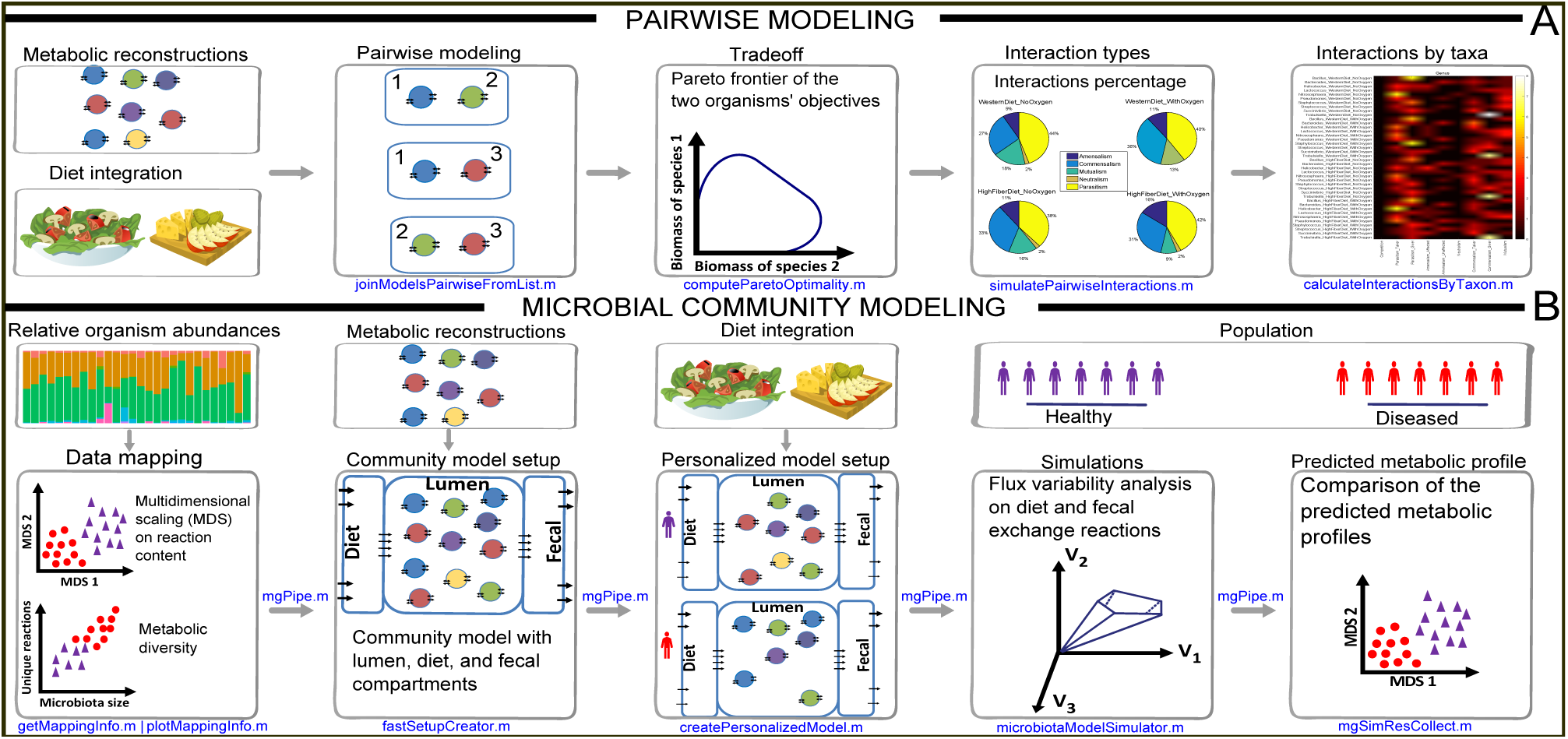
Overview of the functionality of the Microbiome Modeling Toolbox. **A.** Pairwise modeling of microbe-microbe and host-microbe interactions. **B.** Microbial community modeling.

## 2 Features

The Microbiome Modeling Toolbox (Figure 1) enables the generation, simulation, and interpretation of 1. pairwise microbe-microbe and host-microbe interactions, and 2. personalized microbial community models.

### Pairwise interactions

The pairwise interaction analysis determines metabolic exchange between two reconstructions. Therefore, a joint matrix of two individual genome-scale reconstructions is generated, which enables them to freely exchange metabolites (Figure 1A). A uniform nomenclature of reaction and metabolite abbreviations in the reconstructions is required. Defined nutrient input, e.g., a particular medium formulation, can be applied via the shared compartment using the corresponding exchange reactions. The pairwise microbial models can be investigated for six possible interaction types (i.e., competition, parasitism, amensalism, neutralism, commensalism, and mutualism) and Pareto optimality frontiers can be calculated. The toolbox allows for the integrative analysis of any number of reconstructions, including the human metabolic reonstruction [1]. The tutorials *MicrobeMicrobeInteractions* and *HostMicrobeInteractions* provides a detailed overview of the implemented functionalities.

### Microbial community modeling

To allow the analysis of metagenomic data in the context of metabolic modeling, we created a pipeline within the Microbiome Toolbox, *mgPipe* (Figure 1B). This pipeline requires the identified microbial strains, or species, along with relative abundance data for each sample. This information can be obtained from the metagenomic data using bioinformatic tools (e.g., QIIME 2 [2]). *mgPipe* is divided into three parts: 1. the analysis of individuals’ specific microbes abundances, including metabolic diversity and classical multidimensional scaling of the reactions in the identified microbes. 2. Construction of a personalized microbial community model using the identified microbes and their relative abundance data. For each each personalized model, the corresponding microbial reconstructions are joined by adding reactions to each microbial reconstruction transporting metabolites from the extracellular space to the common lumen compartment. Metabolites present in the lumen compartment are connected to a diet and fecal compartment, enabling the uptake and secretion from/to the environment, respectively. We optimized the combination of hundreds of reconstructions using static parallelization. In each microbial community model, the community biomass reaction is personalized using the relative abundance data. Finally, coupling constraints [3] are applied to couple the flux through each microbial reaction to its corresponding biomass reaction flux. And 3. simulation of the personalized microbial community models under different diet regimes, e.g., using flux variability analysis [5]. The differences between maximal uptake and secretion fluxes provide a metabolic profile for each microbial community sample, which can be analyzed using classical multidimensional scaling analysis. Diet-specific constraints (e.g., obtained from http://vmh.life/nutrition) can be applied to the corresponding diet exchange reactions.

## 3 Implementation

The Microbiome Modeling Toolbox is written in MATLAB (Mathworks, Inc.). It is available at https://git.io/microbiomeModelingToolbox together with extensive documentation and tutorials.

## 4 Discussion

The Microbiome Modeling Toolbox enables the user to investigate microbial interactions at a large scale. Moreover, metagenomically-derived data can be integrated with microbial metabolic reconstructions permitting the prediction of altered functional assessment of different microbial communities, e.g., in health and disease.

